# Post-mortem T_2_^*^- weighted MRI imaging of cortical iron reflects severity of Alzheimer’s Disease

**DOI:** 10.1101/279513

**Authors:** Marjolein Bulk, Boyd Kenkhuis, Linda M. van der Graaf, Jelle J. Goeman, Remco Natté, Louise van der Weerd

## Abstract

The value of iron-based MRI changes for the diagnosis and staging of Alzheimer’s disease (AD) depends on an association between cortical iron accumulation and AD pathology. Therefore, this study determined the cortical distribution pattern of MRI contrast changes in cortical regions selected based on the known distribution pattern of tau pathology and investigated whether MRI contrast changes reflect the underlying AD pathology in the different lobes.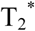-weighted MRI was performed on post-mortem cortical tissue of controls, late-onset AD, and early-onset AD followed by histology and correlation analyses. Combining ex-vivo high-resolution MRI and histopathology revealed that: LOAD and EOAD have a different distribution pattern of AD pathological hallmarks and MRI contrast changes over the cortex, with EOAD showing more severe MRI changes; (2) per lobe, severity of AD pathological hallmarks correlates with iron accumulation, and hence with MRI. Therefore, iron-sensitive MRI sequences allow detection of the cortical distribution pattern of AD pathology ex-vivo.

**Abbreviations:**

AD: Alzheimer’s disease
EOAD: early-onset AD
GM: gray matter
IRP: iron regulating proteins
LOAD: late-onset AD
MCI: mild cognitive impairment
PBS: phosphate buffered saline
QSM: quantitative susceptibility mapping
WM: white matter

## 1. Introduction

The histopathological hallmarks of Alzheimer’s disease (AD) are cerebral amyloid β protein (Aβ) and the paired helical filaments of hyperphosphorylated tau protein. During disease progression, both pathologies distribute throughout the brain in characteristic patterns, which are not the same for Aβ and tau. In end stage AD, as described previously, both Aβ plaques and tau tangles are found throughout the cortex (Alafuzoff et al., 2008; Braak et al., 2006; Braak and Braak, 1991). However, not all cortical regions are equally affected; the temporal lobe is most affected by both pathologies, whereas the somatomotor and stiatal cortex are relatively spared (Braak et al., 2006). The staging of tau pathology as proposed by Braak better reflects cognitive impairment and is therefore considered as a more reliable biomarker for disease progression (Murray et al., 2015).

Although the progression of disease as described by Braak are widely accepted and used, distribution of AD pathology might differ between individuals and certain subtypes of AD (van der Flier et al., 2011). It remains to be proven whether patients with a disease onset before 65 years old, often referred to as early-onset AD (EOAD), and late-onset AD (LOAD) patients show identical distribution patterns. It is already known that autosomal dominant mutations in the PSEN1, PSEN2 or APP gene often results in an earlier disease onset (Alzheimer’s Association, 2016; Reitz and Mayeux, 2014; van der Flier et al., 2011), but EOAD patients also have a more rapid cognitive decline (Koss et al., 1996; van der Vlies et al., 2009) and a more severe post-mortem pathology (Cho et al., 2013; Marshall et al., 2007). In addition, differences in the progression of gray matter (GM) atrophy have been reported, with EOAD patients showing a more widespread atrophy of the cortex, whereas in LOAD patients predominantly the temporal regions are affected (Migliaccio et al., 2015; Moller et al., 2013), suggesting different patterns of atrophy within the AD spectrum.

Apart from amyloid and tau, iron accumulation has received much attention as a putative biomarker and as a modulator of disease progression (Hare et al., 2013; Peters et al., 2015; Ward et al., 2014). Recently, we and others showed that high-field (7T) MRI can be used to detect susceptibility-based cortical contrast changes, which are caused by differences in iron accumulation (Bulk et al., 2018; Nabuurs et al., 2011; van Rooden et al., 2009; van Rooden et al., 2014). Susceptibility differences have been reported between non-demented healthy controls and AD subjects *in vivo* (van Rooden et al., 2014), with EOAD patients showing an increased cortical phase shift, indicating more iron accumulation, compared to LOAD patients in specific cortical regions (van Rooden et al., 2015). The same technique has been used to show that in subjects with subjective cognitive impairment, which is thought to predict a future diagnosis of AD, increased cortical phase shift is associated with a poorer memory performance (van Rooden et al., 2016), and Ayton et al. recently showed that higher hippocampal susceptibility is associated with more rapid cognitive deterioration in MCI and AD patients (Ayton et al., 2017).

The value of iron-based MRI changes for the diagnosis and staging of AD depends on an association between cortical iron accumulation and AD pathology. This association was recently confirmed by a post-mortem study which showed a high correlation between cortical iron accumulation and the amount of Aβ plaques, tau pathology, and Braak score in the frontal cortex (van Duijn et al., 2017). In another study, we showed that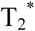-weighted contrast on post-mortem MRI scans correlates with histologically confirmed iron changes. In the frontal cortex, both iron and myelin were found as important contributors to the observed MRI contrast changes, with EOAD patients showing more severe iron accumulation and MRI contrast changes than LOAD patients (Bulk et al., 2018). These previous studies were limited to the frontal cortex, thus the cortical distribution pattern of iron accumulation and the associated MRI contrast changes over the entire cortex remains unknown.

Therefore, the first objective of this study was to determine the distribution pattern of the observed MRI contrast changes in tissue blocks from different cortical regions selected based on the distribution pattern of tau pathology. Secondly, using both MRI and histology, we correlated the MRI contrast changes with the severity of the AD pathology in each region to investigate whether the observed MRI contrast changes reflect the underlying AD pathology in the different lobes. Finally, as mentioned above, differences between EOAD and LOAD patients in distribution patterns of pathology and cortical iron accumulation have been suggested. Therefore, we compared the cortical distribution patterns in both AD subtypes.

## 2. Material and Methods

### 2.1 Study design

To study the cortical distribution pattern of MRI contrast changes in AD, 7T MRI was performed on post-mortem brain samples, using tissue blocks from all four cerebral lobes. The tissue blocks were selected based on the known distribution pattern of tau pathology as described previously by Braak (Braak et al., 2006). MRI contrast changes and histopathology were assessed with the use of semi-quantitative ordinal scoring criteria. Correlation analysis was done to investigate the correlation of MRI contrast changes and severity of AD pathology.

In total, tissue samples of 13 control subjects with no clinical or pathological signs of AD, and 10 LOAD and 11 EOAD patients with a clinical and confirmed pathological diagnosis of AD were included. Control subjects and LOAD patients were sex- and age-matched. LOAD and EOAD patients were matched on sex and Braak scores; by definition EOAD patients had a significant lower age of onset and age of death compared to LOAD patients (p < 0.001). Patient-group characteristics are presented in Table 1.

**Table 1.** Case characteristics.

|  | **Controls (n=13)** | **LOAD (n=10)** | **EOAD (n=11)** |
| --- | --- | --- | --- |
| Mean age of onset, y (range) | - | 79,4 (69-89) | 52,5 (34-64)<sup>++</sup> |
| Mean age of death, y (range) | 80,2 (64-93) | 85,8 (73-96) | 67,3 (43-91) <sup>*,++</sup> |
| Male/female | 5/8 | 4/6 | 4/7 |
| Braak, median (range) | 2 (0-3) | 5 (4-6) ** | 6 (4-6) ** |
| *APOE*- ε4 carriers | 1/8 | 8/10* | 5/11 |
| Mutation (number of known cases) | Unknown | Unknown | APP (1), PSEN1 (1) |
| Post-mortem interval, mean (sd) | 6h:39m (1h:21m) | 5h:17m (1h:20m)* | 5h:01m (1h:02m)* |
| Duration formalin-fixation, median (range) | 1 year (0-3) | 1 year (0-3) | 1 year (0-3) |

\* Indicates a significant difference between controls and LOAD and/or controls and EOAD patients. +

### 2.2 Brain sample preparation

Brain tissue from all subjects was obtained from the Netherlands Brain Bank (NBB, Netherlands Institute for Neuroscience Amsterdam) and the Normal Aging Brain Collection (VU, Amsterdam). Following the Dutch national ethical guidelines, anonymity of all subjects was preserved by using a coded system for the tissue samples. For all subjects, tissue samples were retrieved from specific anatomical locations based on the known distribution pattern of tau pathology as described previously by Braak (Braak et al., 2006), including the middle frontal gyrus, superior parietal gyrus, superior occipital gyrus and middle temporal gyrus. Tissue was fixed in 4% phosphate-buffered formalin and to avoid formalin-induced artefacts on MRI, only material fixed for a minimum of 4 weeks and a maximum of three years was selected (van Duijn et al., 2011). In both control subjects and AD patients, most cases were fixed for not more than one year. No significant differences in duration of formalin fixation were found between the groups.

Tissue blocks of approximately 20×15×15mm were resected and put in a 15 ml tube (Greiner Bio-One). Before MRI, the MR relaxation parameters were partially restored by removing residual formalin and placing the tissue block in phosphate buffered saline (PBS) for > 24 hours (Shepherd et al., 2009). Before scanning, PBS was replaced with a proton-free fluid (Fomblin LC08, Solvay). Care was taken to avoid the inclusion of trapped air bubbles.

### 2.3 Post-mortem MRI acquisition

MRI scans were made on a 7T horizontal bore Bruker MRI system equipped with a 23 mm receiver coil and Paravision 5.1 imaging software (Bruker Biospin, Ettlingen, Germany). Multiple gradient echo scans with a total imaging time of 210 minutes were acquired from each brain sample with repetition time = 75.0 ms, echo times = 12.5, 23.2, 33.9 and 44.6 ms, flip angle = 25° at 100 μm isotropic resolution with 20 signal averages.

### 2.4 Post-mortem MRI analysis

Magnitude images with an echo time of 33.9 ms were selected because these magnitude images visually had the best contrast compared to the other echoes, based also on previous publications (Bulk et al., 2018; van Rooden et al., 2009). Assessment of the cortices was performed using a pre-defined scoring system for assessing MRI contrast changes on 7T MRI in the frontal cortex as described by Bulk et al (Bulk et al., 2018). These included assessments of cortical homogeneity and the presence of a diffuse hypointense band.

A normal cortex was defined as either a cortex containing one homogeneous hyperintense layer in comparison to the adjacent white matter (WM), or a homogeneous cortex containing two or three well-delineated layers observed as normal cortical lamination. The superficial layer has a higher signal intensity compared with the deepest layers. Additionally, in a cortex with three layers a thin third layer with a lower signal intensity compared with the other layers as described previously in cortical regions with highly myelinated layers could be observed. This thin layer is known as the line of Baillarger, or in the striate cortex specifically as the line of Gennari (Braak et al., 1989). In a normal cortex with normal cortical lamination, no diffuse hypointense band was observed (Fig. 1, thick arrow). An abnormal cortex was defined as a cortex with a granular/patchy structure containing foci of signal loss. In addition, a diffuse hypointense band was observed as diffuse areas of lower signal intensity covering the middle cortical layers, obscuring normal cortical lamination (Fig. 1, arrow).

**Figure 1.**
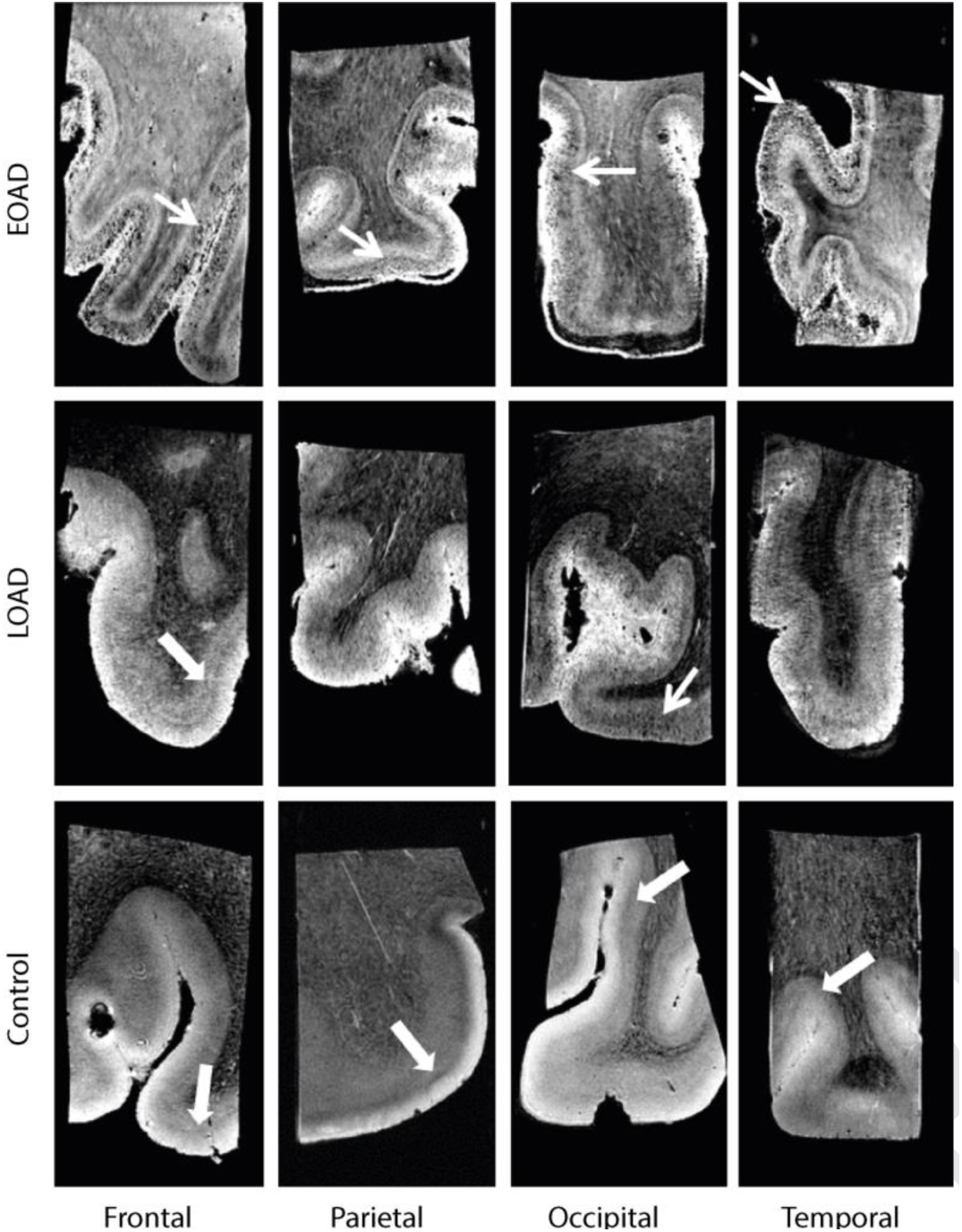
Representative MRI images of a control, LOAD and EOAD subject. Control subjects were characterized by a normal cortex defined as a homogeneous cortex including normal cortical lamination (thick arrow). AD patients have a different cortical appearance on 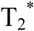-weighted MRI compared to control subjects, characterized by an inhomogeneous cortex with a granular and/or patchy structure and foci of signal loss (LOAD and EOAD subject) and a diffuse hypointense band covering and obscuring the central layers of the cortex (arrow).

All images were scored for absence or presence of above-mentioned criteria. As previously explained (Bulk et al., 2018), different scoring values were used per criteria to indicate presence, partially presence or absence of the criteria. A weakly or intermittently observed diffuse hypointense band throughout the cortical ribbon was scored as partially present. A total score was calculated from the values of each criteria, with a minimum score of 0/4 indicating a normal cortex and a maximum of 4/4 indicating an abnormal cortex. Window settings were standardized for each image and all scans were scored by two independent blinded trained observers (MB, BK). Consensus was reached in all cases of disagreement.

Lastly, to indicate severity of the MRI contrast changes as used in Fig. 5 a severity score was calculated based on the presence of a hypointense diffuse band on MRI. The scoring values for the presence of a diffuse hypointense band (0,1,2) were multiplied by the frequency of each criteria resulting in a sum score for each cortical region. Severity ranges from 0 to 22, with 0 indicating low severity and 20 high severity.

**Figure 5.**
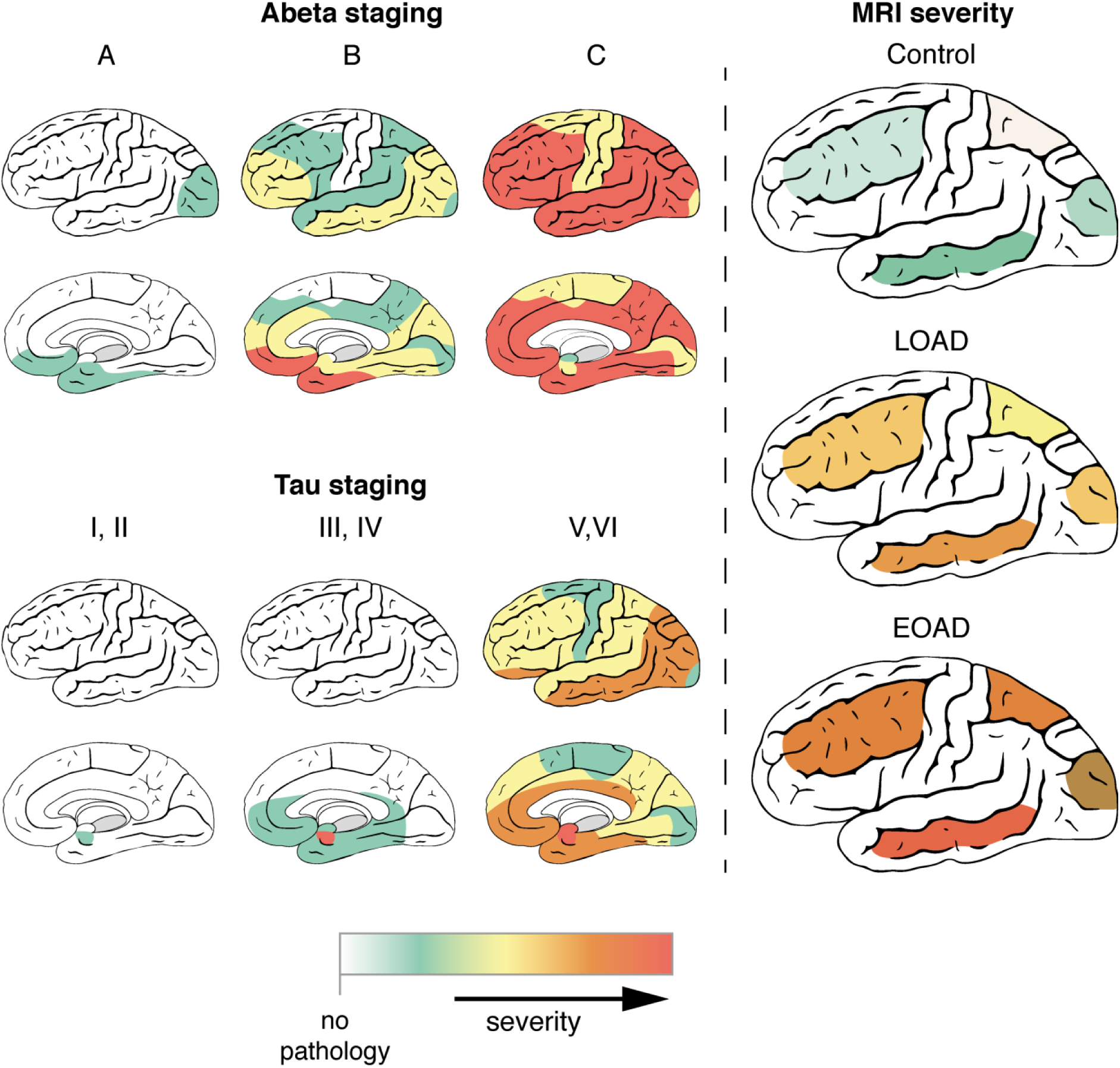
Distribution of AD pathology and MRI contrast changes over the cortex. During disease progression, both AD pathological hallmarks spread throughout the brain in characteristic patterns, which are not the same for Aβ and tau. As described previously, Aβ plaques are first found in the basal parts of the frontal, occipital, and temporal lobe and gradually spread throughout the whole cortex. In later stages, Aβ plaques are also found in allocortical regions, the striatum, brainstem nuclei and eventually in the cerebellum. In contrast to Aβ plaques, tau tangles are first detected in the transentorhinal region of the middle temporal lobe. Subsequently, the basal parts of the frontal and temporal lobe are affected and finally the entire cortex, including the striatal cortex. In the histologically determined stages of AD pathology as well as the MRI contrast changes, the temporal lobe is the most affected region corresponding to the distribution pattern of tau pathology. Even in controls, which have only limited tau pathology and MRI contrast changes, the temporal lobe is most affected. Severity scales ranges for abeta and tau staging from no pathology (white) to severe pathology (red). For MRI, MRI severity scores were calculated for specific cortical regions based on presence of a diffuse hypointense band (Methods 2.4). MRI severity scales ranges from severity not determined (white) to severe MRI changes (red).

### 2.5.Histology and immunohistochemistry

The same tissue blocks as used for MRI, were also used for histology. Tissue blocks were embedded in paraffin and serially cut into 8-μm and 20-μm-thick sections. Histochemical iron detection was done as previously described (Bulk et al., 2018; van Duijn et al., 2013). After deparaffinization, 20-μm-thick sections were incubated for 80 minutes in 1% potassium ferrocyanide, washed, followed by 100 minutes incubation in methanol with 0.01 M NaN3 and 0.3% H2O2. Subsequently, sections were washed with 0.1 M phosphate buffer followed by 80 minutes incubation in a solution containing 0.025% 3‘3-diaminobenzidine-tetrahydrochloride (DAB, (DakoCytomation)) and 0.005% H2O2 in 0.1 M phosphate buffer. The reaction was stopped by washing with tap water.

Another 20-μm-thick section was used for immunohistochemical detection of myelin, and the 8-μm sections were stained for Aβ and paired helical filament-tau (Supplementary Table 1). All sections were treated with 0.3% H2O2 in methanol to block endogenous peroxidase activity. Depending on the primary antibody this step was followed by an antigen retrieval step. Primary antibodies were incubated overnight at room temperature. The secondary antibody was incubated for one hour followed by a 30 minutes incubation with avidin-biotin complex (ABC, Vector Labs, CA, USA). Signal enhancement was completed by immersion in DAB. The 8-μm sections were counterstained with Harris Haematoxylin. All slides were digitized using an automatic bright field microscope (Philips Ultra Fast Scanner, Philips, Netherlands) for microscopic evaluation.

### 2.6.Semi-quantitative scoring of pathology

Histopathological stainings have been performed for: Aβ, tau tangles, iron and myelin. The same stainings have previously been investigated (Bulk et al., 2018; van Duijn et al., 2013), and the same semi-quantitative scoring system was used in this study. All substrates were assessed by two trained blinded observers (MB, BK). In case of disagreement or doubtful scores, the specific subject was assessed by a neuropathologist (RN).

In short, severity of cortical Aβ plaque load was divided into four categories: absent (no Aβ plaques [scoring value 0]), mild (one or scattered Aβ plaques present [scoring value 1]), moderate (widespread Aβ plaques, but with one or more areas in layer II-IV without or with only little amount of Aβ plaques [scoring value 2]), severe (diffuse high Aβ plaque density in layer II-IV [scoring value 3]).

For hyperphosporylated tau, the following categories were defined: absent (no hyperphosphorylated tau labelling present [scoring value 0]), light (occasional hyper-phosphorylated tau labelling of individual cells or neuropil threads [scoring value 1]), mild (at least one 200x area with hyper-phosphorylated tau labelling, as adapted from figure 5B in Alafuzoff et al. (Alafuzoff et al., 2008) [scoring value 2]), moderate (macroscopically visible diffuse hyperphosphorylated tau labeling of part of the cortex, or restricted to one cortical layer [scoring value 3]), severe (macroscopically visible diffuse hyperphosphorylated tau labelling of more than one visible layer (mostly layer V and II/III) for the complete cortical ribbon [scoring value 4]).

Cortical iron and myelin were classified as no cortical staining (scoring value 1)), normal cortical staining (scoring value 2), partially thickened cortical staining (scoring value 3) or abnormal, diffuse band shaped, cortical staining [scoring value 4]. Abnormal cortical iron and myelin were both defined as a diffuse band-shaped pattern of respectively increased iron labelling and increased myelin immunohistochemistry, covering the middle cortical layers (III/IV), sometimes extending to layer II and V. A weak or intermittently observed band-shaped staining intensity throughout the cortical ribbon was scored as partially thickened cortical staining.

### 2.7. Statistical methods

Continuous variables, as for example age, were compared between groups using a one-way ANOVA. A chi-square test was used to compare categorical variables between groups and between regions. Since three groups were compared post-hoc chi-square and anova tests were done using the method of Shaffer (Shaffer, 1986), that allows post hoc tests at the full level. Semi-quantitative MRI scores were correlated to semi-quantitative scores for Aβ plaque load, tau load, cortical iron accumulation and cortical myelin changes using a linear by linear association test for ordinal variables. All statistical analyses were performed using Statistical Package of Social Sciences (SPSS, version 23; SPSS, Chicago, USA). A significance level of 0.05 was used.

## 3. Results

### 3.1.Cortical distribution pattern of MRI contrast changes

In a previous ex-vivo 7T imaging study, we showed that AD patients have a different cortical appearance on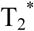-weighted MRI compared to control subjects, characterized by inhomogeneities and a diffuse hypointense band covering the central layers of the frontal cortex. In the current study, we determined the cortical distribution pattern of MRI contrast changes over the brain, using tissue blocks of the frontal, parietal, occipital, and temporal cortex and the previously used MRI scoring system (Fig. 1) (Bulk et al., 2018).

Both an inhomogeneous cortex and a diffuse hypointense band were more frequently found in LOAD and EOAD patients compared to control subjects (Table 2). Whereas an inhomogeneous cortex was observed equally across all cortical regions in AD patients, regardless of age of onset (Fig. 2A), the presence of a diffuse hypointense band showed a cortical distribution pattern (Fig. 2B). Generally in LOAD patients we observed the diffuse hypointense band more prominently in the temporal and occipital regions, whereas the frontal regions were less severely affected. In contrast, in the EOAD patients we frequently observed heavy involvement of all lobes. Furthermore, the diffuse hypointense band was significantly more frequently visible in EOAD patients than in LOAD patients (p<0.05, Table 2, Fig. 2B).

**Table 2.** MRI and histopathological findings

|  | Controls (n=13) | LOAD (n=10) | EOAD (n=11) |
| --- | --- | --- | --- |
| **Homogeneity of the cortex, homogeneous/inhomogeneous** |  |  |  |
| Frontal middle gyrus | 9/4 | 0/10* | 2/9* |
| Parietal superior gyrus | 11/2 | 1/9** | 3/8* |
| Occipital superior gyrus | 10/3 | 0/10** | 2/9* |
| Temporal middle gyrus | 10/3 | 1/9* | 1/10* |
| **Diffuse hypointense band, absent/partially present/present** |  |  |  |
| Frontal middle gyrus | 11/2/0 | 0/8/2** | 2/3/6*,+ |
| Parietal superior gyrus | 12/1/0 | 1/7/2** | 3/1/7*,+ |
| Occipital superior gyrus | 10/3/0 | 1/6/3** | 2/3/5* |
| Temporal middle gyrus | 8/5/0 | 1/4/5** | 1/2/8* |
| **Total MRI score, mean (sd)** |  |  |  |
| Frontal middle gyrus | 0.77 (1.092) | 3.20 (0.422)** | 3.00 (1.549)** |
| Parietal superior gyrus | 0.38 (0.961) | 2.90 (1.101)** | 2.82 (1.834)** |
| Occipital superior gyrus | 0.69 (1.316) | 3.20 (0.632)** | 3.00 (1.265)** |
| Temporal middle gyrus | 0.85 (1.280) | 3.20 (1.229)** | 3.45 (1.214)** |
| **Aβ load, median (range)** |  |  |  |
| Frontal middle gyrus | 1 (0-2) | 3 (2-3)* | 3 (-)**+ |
| Parietal superior gyrus | 0 (0-2) | 3 (2-3)** | 3 (-)** |
| Occipital superior gyrus | 0 (0-2) | 3 (2-3)** | 3 (2-3)** |
| Temporal middle gyrus | 0 (0-2) | 3 (1-3)** | 3 (-)** |
| **Tau load, median (range)** |  |  |  |
| Frontal middle gyrus | 0 (0-1) | 2 (1-3)** | 4 (3-4)**+ |
| Parietal superior gyrus | 0 (0-1) | 3 (1-4)** | 4 (3-4)**+ |
| Occipital superior gyrus | 0 (0-1) | 3 (0-3)* | 3 (3-4)** |
| Temporal middle gyrus | 1 (0-1)# | 3 (2-4)* | 4 (3-4)** |
| **Cortical iron accumulation, median (range)** |  |  |  |
| Frontal middle gyrus | 2 (1-2) | 2 (1-4) | 3 (1-4)* |
| Parietal superior gyrus | 2 (1-2) | 2.5 (1-4) | 3 (3-4)** |
| Occipital superior gyrus | 2 (1-2) | 2 (2-3) | 3 (2-4)* |
| Temporal middle gyrus | 1.5 (1-2) | 2.5 (1-4) | 4 (1-4)* |
| **Cortical myelin organization, median (range)** |  |  |  |
| Frontal middle gyrus | 2 (-) | 2 (2-3) | 4 (2-4)*+ |
| Parietal superior gyrus | 2 (2-3) | 2 (2-3) | 4 (3-4)**+ |
| Occipital superior gyrus | 2 (-) | 2 (2-3)* | 4 (2-4)**+ |
| Temporal middle gyrus | 2 (-) | 2 (2-3)* | 4 (2-4)*+ |

\* Indicates a significant difference between controls and LOAD and/or controls and EOAD patients. + Indicates a significant difference between LOAD and EOAD patients. # Indicates a significant differences between cortical regions within the control group. \*/+/# indicates $p < 0.05$ and \*\*/+ indicates $p < 0.001$ .

**Figure 2.**
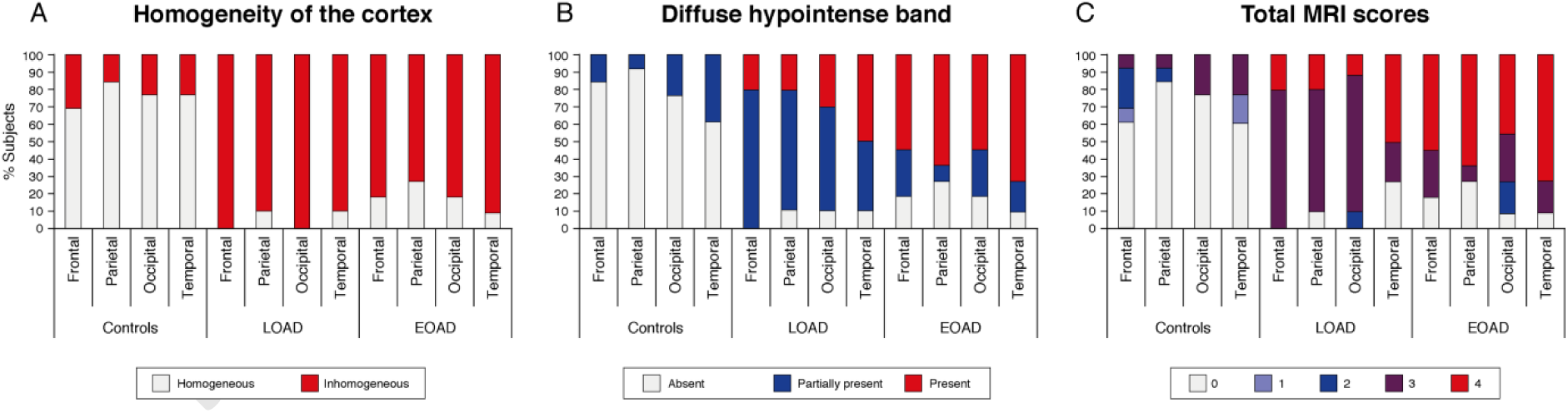
Spreading of MRI contrast changes over the cortex. **(A)** An inhomogeneous cortex was observed equally across all cortical regions in AD patients regardless of age of onset. (B) The presence of a diffuse hypointense band showed a cortical spreading pattern. In LOAD patients, the diffuse hypointense band was most frequently observed in the temporal and occipital regions, whereas the frontal regions were less severely affected. In contrast, in the EOAD patients all regions were similarly affected, and the diffuse hypointense band was significantly more visible in LOAD patients than in EOAD patients. (C) For all cortical regions, control subjects had significantly lower total MRI scores compared to both AD groups indicating a normal cortical appearance on MRI. The temporal cortex was the most affected region in LOAD patients, followed by the other cortical regions. In EOAD patients all cortical regions were heavily affected.

By summation of the individual scoring criteria, a total MRI score was calculated per cortical region. A low score indicated a normally-appearing cortex and a high score an abnormally-appearing cortex. For all cortical regions, control subjects had significantly lower total MRI scores compared to both AD groups (Table 2, Fig. 2C).

### 3.2 Cortical distribution pattern of iron, myelin and AD pathology

To assess the correlation between the pathological burden and the distribution pattern of AD pathology over the different cortical regions, the same tissue blocks as used for MRI were processed for histological analysis. Aβ plaques were abundantly present in all AD patients and all cortical regions were heavily affected. As expected, Aβ plaques were also present in some control subjects (Fig. 3A). Following Braak staging, the distribution pattern of hyperphosphorylated tau showed a predominance for the temporal cortex, with 69% of the control subjects already showing some presence of tau and nearly all LOAD patients showing severe tau pathology. In contrast, in EOAD patients all cortical regions were equally affected. In addition, EOAD patients showed significantly more hyperphosphorylated tau in the frontal and parietal cortex compared to LOAD patients (Table 2, Fig. 3B).

**Figure 3.**
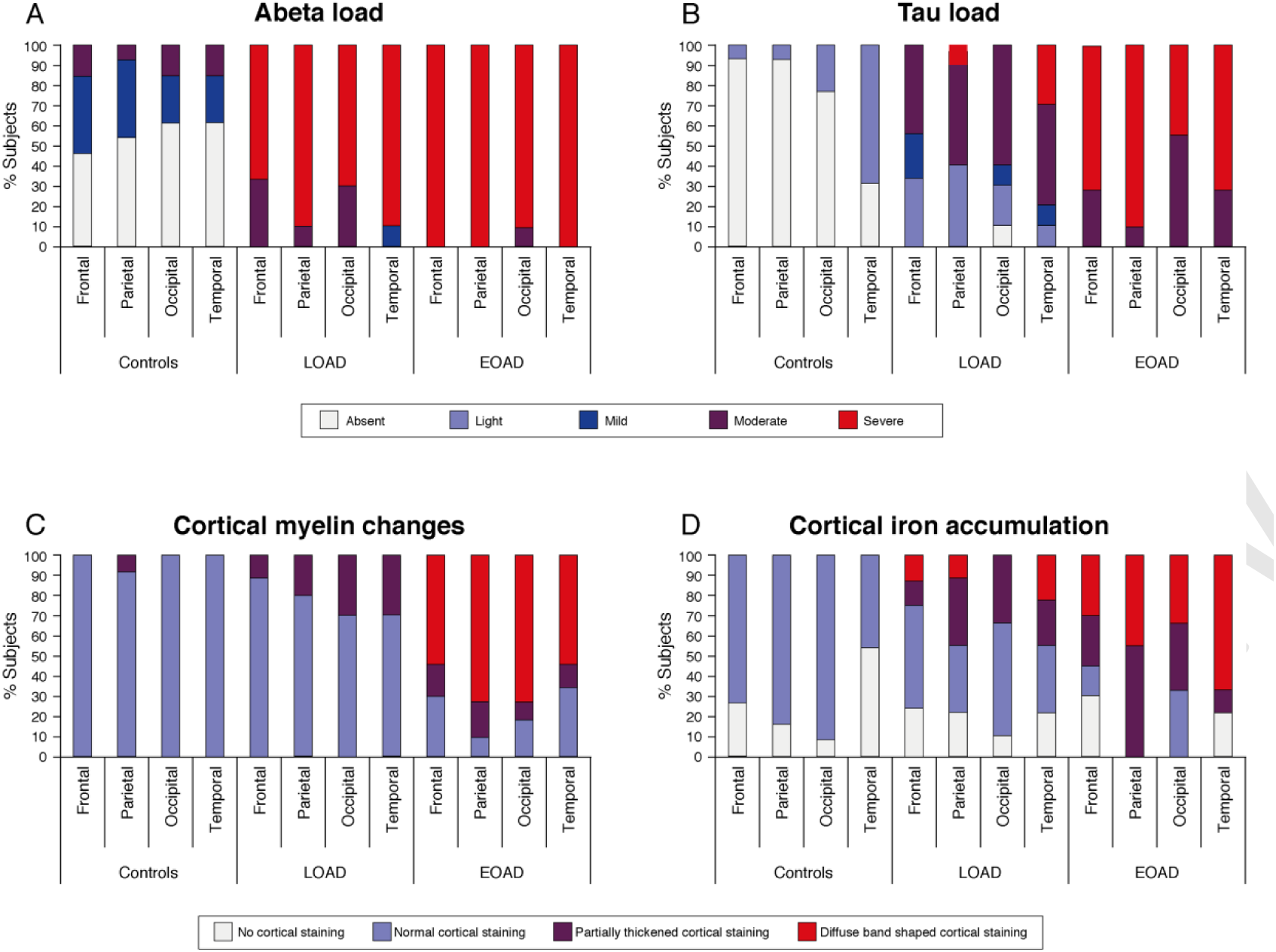
Spreading pattern of AD pathology and cortical iron and myelin changes over the cortex. (A) Aβ plaques were abundantly present in all AD patients and all cortical regions were heavily affected. (B) The spreading pattern of hyperphosphorylated tau showed a predominance for the temporal cortex, with 69% of the control subjects already showing some presence of tau and nearly all LOAD patients showing severe tau pathology. In EOAD patients all cortical regions were equally affected. (C) Nearly all control subjects and the majority of the LOAD patients showed a normal myelin distribution. In contrast, the majority of the EOAD patients showed an abnormal appearance of cortical myelin across all cortical regions. (D) Compared to control subjects, in whom normal myelin-associated iron was observed, some AD patients showed an altered pattern of diffuse myelin-associated iron. In both LOAD and EOAD patients, the temporal cortex showed to most severe iron accumulation compared to the other cortical regions.

Since we previously showed that changes in cortical iron and myelin organization are the predominant sources of the observed MRI contrast changes in AD patients, cortical myelin and iron were also investigated (Table 2, Fig. 3C,D). Nearly all control subjects and the majority of the LOAD patients showed a normal myelin distribution with an increased staining intensity in myelin-rich areas (WM, lines of Baillarger and myelinated fiber bundles transversing from the WM into the GM). In contrast, the majority of the EOAD patients showed an abnormal appearance of cortical myelin, observed as a diffuse band-shaped increased myelin staining of the middle layers of the cortex (III/IV), sometimes extending to layer II and V. This diffuse band-shaped increased myelin staining intensity was significantly more observed in EOAD patients compared to controls and LOAD patients across all cortical regions.

Compared to all control subjects, in whom normal myelin-associated iron was observed (i.e. in myelin-rich areas), some AD patients showed an altered pattern of cortical iron. This was observed as a diffuse band-shaped pattern of increased iron associated with and without cortical myelin covering the middle layers of the cortex (III/IV), sometimes extending till layer II and V. In both LOAD and EOAD patients, the temporal cortex showed to most severe iron accumulation compared to the other cortical regions (Fig. 3D).

### 3.3.MRI contrast is correlated with iron, myelin and AD pathology severity

Since the presence of the diffuse hypointense band on MRI is the most discriminating scoring criterion between control subjects, LOAD and EOAD patients, this scoring criterion was used to investigate the correlation between MRI contrast changes and severity of AD pathology (Fig. 4).

**Figure 4.**
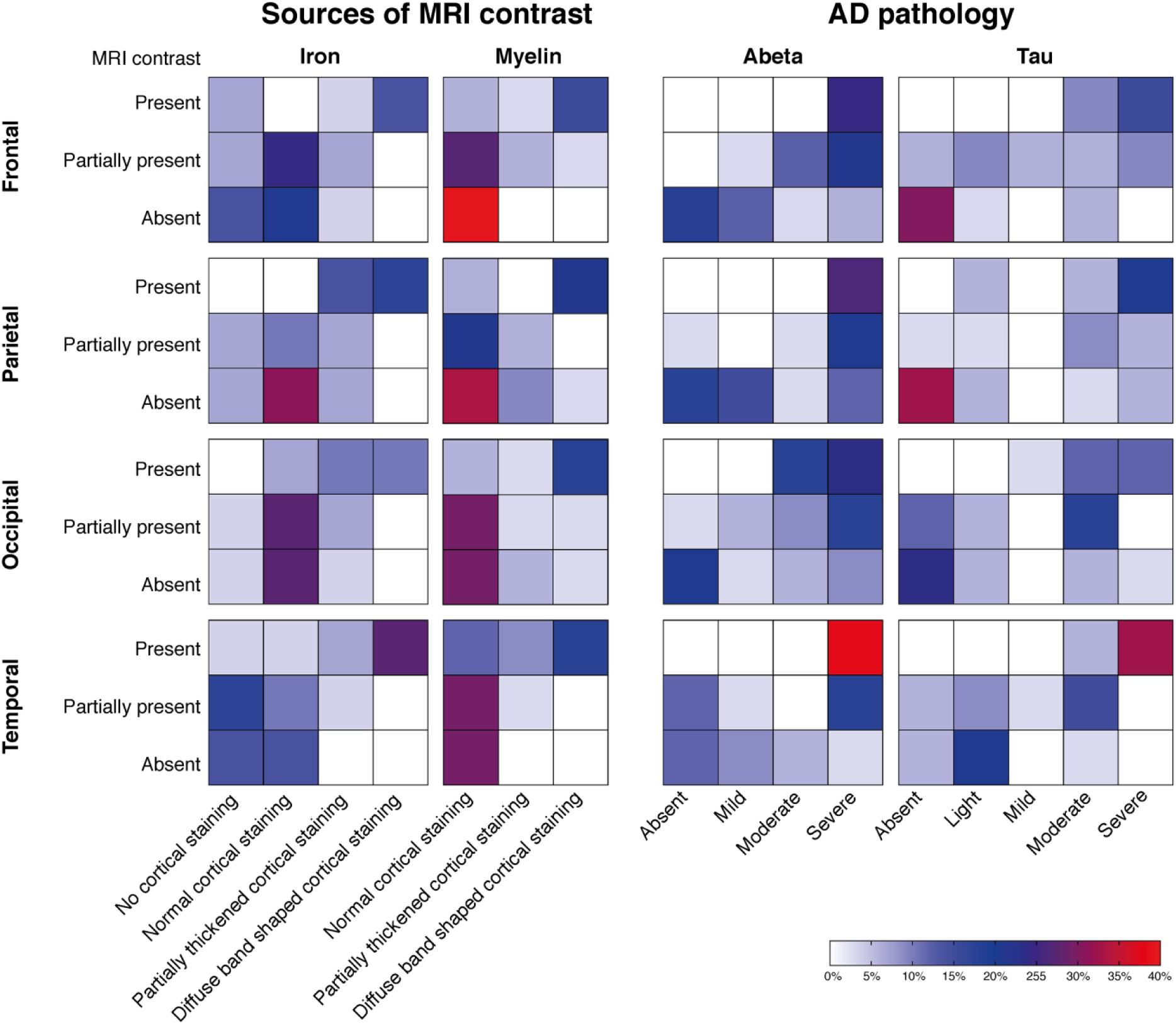
Heatmaps of the correlation between MRI contrast changes and AD pathology severity. The presence of a diffuse hypointense band on MRI was significantly correlated with the semi-quantitative scored amount of cortical iron accumulation, cortical myelin changes, Aβ plaque load and tau load in all cortical regions (P<0.05). In our cohort, a diffuse hypointense band on MRI was always accompanied by moderate-to-high amounts of Aβ plaques and light-to-severe tau load. In subjects without a diffuse hypointense band on MRI, mostly absent-to-low amounts of hyperphosphorylated tau were found.

The presence of a diffuse hypointense band on MRI was significantly correlated with the semi-quantitative scored amount of cortical iron accumulation, Aβ plaque load, tau load, and cortical myelin changes in all cortical regions (P<0.05). In our cohort, a diffuse hypointense band on MRI was always accompanied by moderate-to-high amounts of Aβ plaques and light-to-severe tau load. In subjects without a diffuse hypointense band on MRI, mostly absent-to-low amounts of hyperphosphorylated tau were found.

### 3.4.Cortical distribution pattern is not affected by APOE genotype

Besides differences between LOAD and EOAD patients, we also investigated the additional effect of APOE genotype on cortical distribution patterns. However, no difference was found between APOE ε4 carriers and non-carriers on cortical appearance on MRI, pathological burden or cortical distribution pattern of MRI contrast changes and AD pathology (data not shown).

## 4.Discussion

This study aimed to determine whether the cortical distribution pattern of 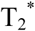-weighted MRI contrast changes reflects the known progression of AD pathology over the cortex. In addition, we investigated whether the MRI contrast changes reflect the severity of AD pathology in the different lobes and described the differences between LOAD and EOAD patients in distribution patterns of both MRI contrast changes and pathology. The combination of ex-vivo high-resolution MRI and histopathology on the same tissue block revealed that: (1) LOAD and EOAD patients have a different distribution pattern of AD pathological hallmarks and MRI contrast changes over the cortex, with EOAD patients showing more severe MRI changes; (2) per lobe, severity of AD pathological hallmarks correlates with iron accumulation, and hence with MRI. As visualized in Fig. 5, in the histologically determined stages of AD pathology as well as the MRI contrast changes, the temporal lobe is the most affected region corresponding to the distribution pattern of tau pathology. Even in controls, which have only limited tau pathology and MRI contrast changes, the temporal lobe is most affected. Therefore, using iron-sensitive MRI sequences, such as 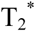-weighted MRI, we can non-invasively detect the cortical distribution pattern of AD pathology ex-vivo.

We observed an abnormal appearance of the cortex in AD patients characterized by inhomogeneities and a diffuse hypointense band. Iron and myelin have been previously reported as an important source for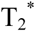-weighted MRI contrast in the cortex (Fukunaga et al., 2010; Langkammer et al., 2010; Stuber et al., 2014; Wallace et al., 2016). We previously showed that also in AD, 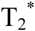-weighted contrast spatially correlated with changes in cortical iron accumulation and myelin organization (Bulk et al., 2018). Even though we did not find a spatial correlation of MRI contrast with the AD pathological hallmarks Aβ and hyperphosphorylated tau on a voxel-by-voxel basis, AD patients could clearly be distinguished from controls, indicating that susceptibility-based contrast changes reflect cortical alterations in AD other than Aβ and tau. In the current study, we did not investigate voxel-by-voxel correlations between MRI and AD pathological hallmarks; instead we focused on correlations of the observed MRI contrast changes with the semi-quantitative scored amount of cortical iron accumulation, Aβ plaque load, tau load, and cortical myelin changes in the different cortical regions.

The significant correlations between MRI and histological scores suggests that cortical iron accumulation, and consequently the MRI contrast, reflects the severity of AD pathology in each lobe. These findings correspond with recent findings on post-mortem AD tissue showing that the degree of iron accumulation is positively correlated with the amount of Aβ plaques and tau load in the frontal cortex (van Duijn et al., 2017). Moreover, an in-vivo study by van Bergen et al., also found increased cortical iron levels measured by quantitative susceptibility mapping (QSM) MRI in PiB-positive APOE-e4 carriers with mild cognitive impairment (van Bergen et al., 2016), indicating an association between cortical iron and presence of Aβ plaques. The correlation between iron imaging and tau pathology has been less studied. Nevertheless, one study using a mouse model of tauopathy found magnetic susceptibility differences in the corpus callosum, striatum, hippocampus and thalamus. Interestingly, these regions exhibited low NFT burden, but increased markers for reactive microglia and astrocytes, again confirming the notion that iron accumulation correlates with AD pathology, but is not directly caused by iron depositing in amyloid and/or tau aggregates (O’Callaghan et al., 2017).

We found a different distribution pattern of AD pathology and MRI contrast changes in LOAD and EOAD patients. Overall, on both MRI and histology LOAD patients followed the typical distribution patterns as described by the Braak stages, with predominantly temporal lobe involvement. In contrast, EOAD patients were generally more affected on both MRI and histology and besides the temporal lobe also the frontal, parietal, and occipital lobes were frequently affected. However, Aβ plaques were abundantly present in all AD patients and had reached a plateau for all cortical regions. The finding of a larger tau burden in EOAD patients is consistent with previous histological studies showing larger amounts of neurofibrillary tangles in EOAD patients (Marshall et al., 2007). Also, the changes in cortical iron accumulation were more frequently observed in EOAD patients. This is consistent with a previous *in vivo* 7T MRI study showing higher cortical phase MRI changes in EOAD compared to LOAD patients, reflecting increased cortical iron accumulation in EOAD (van Rooden et al., 2015).

Our observation of significant involvement of the frontal, parietal and occipital lobe in addition to the temporal lobe in EOAD patients is supported by previous studies. Atrophy for example, is in EOAD patients more widespread, with more prominent involvement of regions other than the temporal lobe as the posterior and frontoparietal areas (Frisoni et al., 2007; Migliaccio et al., 2015). Apart from structural and pathological differences, it is known that LOAD and EOAD patients differ in their clinical presentation. LOAD patients typically present with memory impairment corresponding with both the typical pattern of GM atrophy and our MRI contrast changes in the medial temporal lobe. In contrast, EOAD patients frequently present with atypical symptoms including visual dysfunction, apraxia and dyscalculia while the memory is relatively preserved (Koedam et al., 2010). This atypical presentation seemingly corresponds to the more widespread pattern of GM atrophy and the observed widespread cortical MRI contrast changes.

Apart from differences in iron accumulation due to age of onset, also APOE gene status is linked to iron (Ayton et al., 2015; van Bergen et al., 2016). Ayton et al showed recently that ferritin is strongly associated with CSF ApoE levels. Moreover, ferritin levels were elevated in APOE-e4 carriers (Ayton et al., 2015). The same association was shown by van Bergen et al., in mild cognitive impairment subjects; APOE-e4 carriers showed significantly higher magnetic susceptibility values compared to non-carriers (van Bergen et al., 2016). These studies suggest that the APOE-e4 allele increases the risk of developing AD via iron accumulation in the brain. However, in our study we did not find significant differences between APOE-e4 carriers and non-carriers, most likely due to the small group sizes.

Despite the correlation between iron,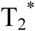-weighted MRI and AD pathology as shown in this study, the temporal correlation of iron accumulation and AD pathology remains to be elucidated. Currently, it is unknown whether iron follows the pathology, is accumulating at the same time, or is even earlier than Aβ and tau. However, in AD and other neurodegenerative diseases iron accumulation have been shown to play an important role in the progression of neurodegeneration (Ayton et al., 2017; van Bergen et al., 2016). Recently it was shown that cortical iron accumulation might accelerate clinical progression in AD in Aβ-positive MCI patients (Ayton et al., 2017). Several mechanisms have been reported for the interaction of iron with amyloid and tau pathology. Apart from the production of ROS as being a well-known consequence of iron accumulation (Zecca et al., 2004), iron also modulates APP cleavage by furin and through the interaction with iron regulating proteins (IRP). Excessive amounts of iron decrease the activity of furin and thereby favour the amyloidgenic pathway of APP cleavage (Silvestri et al., 2008; Ward et al., 2014). In addition, iron might modulate APP translation through IRP; increased amounts of iron might up-regulate APP translation resulting in an increased amount of APP available to be cleaved by the amyloidgenic pathway (Silvestri et al., 2008; Ward et al., 2014). Interestingly, intracellular iron storage and clearance is also regulated by APP and furin, and affected by the other pathological hallmark of AD, namely hyperphosphorylated tau (Hare et al., 2013; Silvestri et al., 2008). As has been shown in-vitro, loss of functional tau disturbs APP-mediated iron export by decreasing trafficking of APP to the neuronal surface, eventually resulting in increased levels of intracellular iron (Lei et al., 2012). Thus, cortical iron accumulation is likely a disease modifier and may be an independent biomarker to predict clinical progression.

## Acknowledgements

The authors thank I. M. Hegeman-Kleinn for her technical assistance and Gerrit Kracht for making the figures. This work was supported by a project grant from the EU Seventh Frame-work Programme: FP7-PEOPLE-2013-IAPP (612360 e BRAINPATH).

## Supplementary Table

**Suppl. Table 1.**
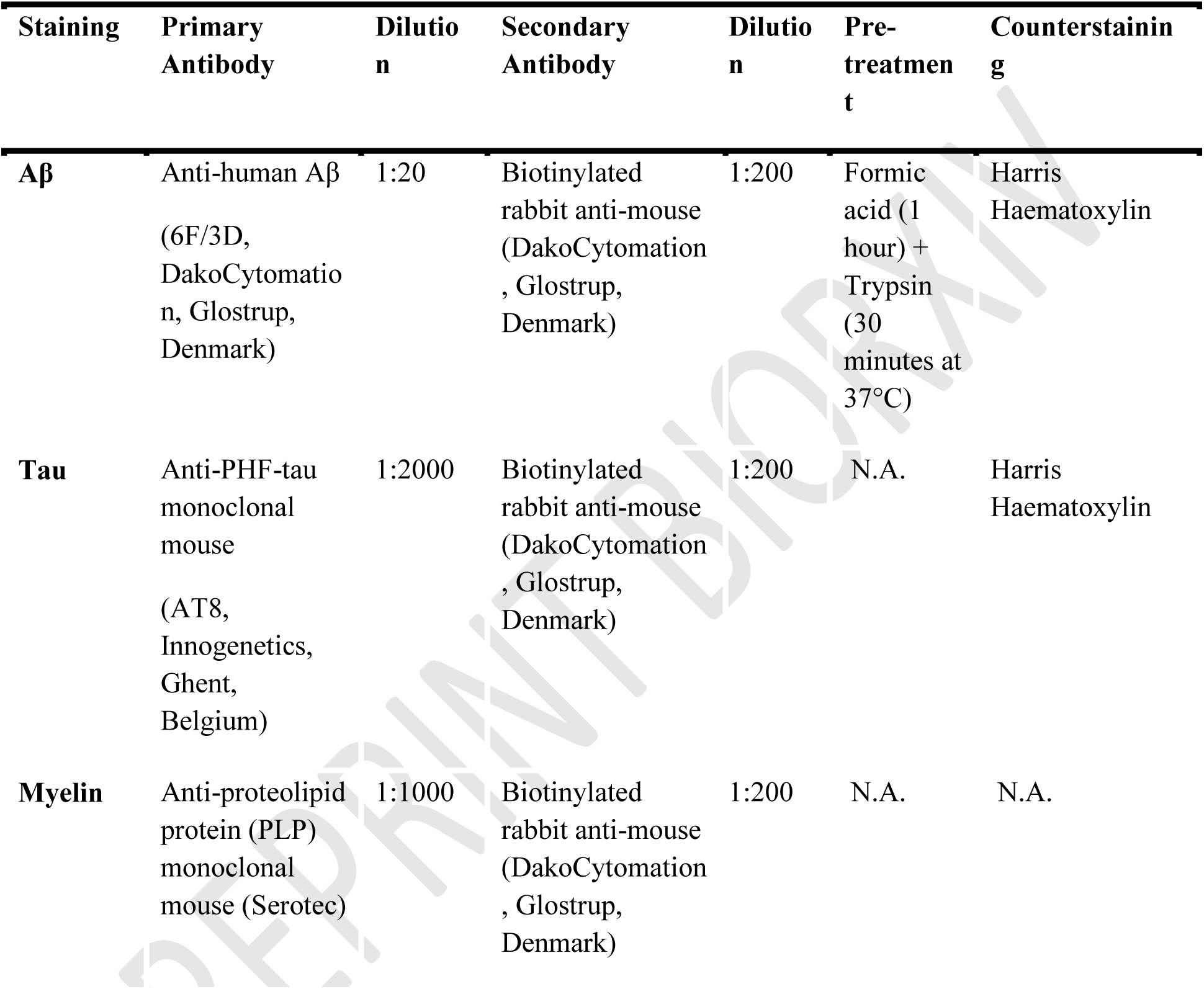
Immunohistochemistry details including primary and secondary antibodies and additional pre-treatments.

| Staining | Primary Antibody | Dilution | Secondary Antibody | Dilution | Pre-treatment | Counterstaining |
| --- | --- | --- | --- | --- | --- | --- |
| **Aβ** | Anti-human Aβ (6F/3D, DakoCytomation, Glostrup, Denmark) | 1:20 | Biotinylated rabbit anti-mouse (DakoCytomation, Glostrup, Denmark) | 1:200 | Formic acid (1 hour) + Trypsin (30 minutes at 37°C) | Harris Haematoxylin |
| **Tau** | Anti-PHF-tau monoclonal mouse (AT8, Innogenetics, Ghent, Belgium) | 1:2000 | Biotinylated rabbit anti-mouse (DakoCytomation, Glostrup, Denmark) | 1:200 | N.A. | Harris Haematoxylin |
| **Myelin** | Anti-proteolipid protein (PLP) monoclonal mouse (Serotec) | 1:1000 | Biotinylated rabbit anti-mouse (DakoCytomation, Glostrup, Denmark) | 1:200 | N.A. | N.A. |

